# A systems approach to the characterization and classification of T-cell responses

**DOI:** 10.1101/175620

**Authors:** Shinobu Yamamoto, Elizabeth Whalen, Daisuke Chujo, Durgha Nattamai, Nicole Baldwin, Kimberly O’Brien, Quynh-Anh Nguyen, Vivian Gersuk, Esperanza Anguiano, Junbao Yang, William W Kwok, Jacques Banchereau, Hideki Ueno, Damien Chaussabel

## Abstract

Types of T-cell responses are categorized on the basis of a limited number of molecular markers selected using *a priori* knowledge about T-cell immunobiology. We sought to develop a novel systems-based approach for the creation of an unbiased framework enabling assessment of antigenic-peptide specific T-cell responses *in vitro*. A meta-analysis of transcriptome data from PBMCs stimulated with a wide range of peptides identified patterns of gene regulation that provided an unbiased classification of types of antigen-specific responses. Further analysis yielded new insight about the molecular processes engaged following antigenic stimulation. This led for instance to the identification of transcription factors not previously studied in the context of T-cell differentiation. Taken together this profiling approach can serve as a basis for the unbiased characterization of antigen-specific responses and as a foundation for the development of novel systems-based immune profiling assays.

## Introduction

T-cells develop in the thymus where they undergo positive and negative selection through which unreactive and auto-reactive T-cells are removed from the lymphocyte pool. Upon antigen encounter with appropriate signals, naïve T-cells further develop into effector and memory T-cells. T-cell fate is influenced by the quantity of antigen and duration of antigen exposure, strength of T-cell receptor (TCR) interaction with peptide-MHC complex, and co-stimulatory signals as well as cytokine environment. However, plasticity of the T-cells is maintained even after they develop into various effector T-cells (1).

Effector CD4^+^ T-cells are central organizers of adaptive immunity, and their current accepted classification includes Th1, Th2, Th9, Th17, Th22, regulatory T (Treg), and follicular helper T (Tfh) cells. Th1 cells induce cell-mediated immunity against intracellular microbes, and are characterized by expression of the transcription factor T-bet and production of effector cytokine IFNG. Th2 cells play a role in allergic inflammation and promote humoral immunity against extracellular microbes, express GATA3, and produce IL4, IL5, and IL13 (2). Th9 cells play roles in allergic and autoimmune inflammations and anti-tumor immunity, express PU.1 and IRF4, and produce IL9 (3). Th17 cells are involved in protection at mucocutaneous sites, express RORγt, and produce IL17, IL21, IL22, and IL26 (4). Th22 cells function in barrier immunity, express AHR, and produce IL22 (5). Tregs express Foxp3, and keep immune responses in check by suppressing the responses partly by secretion of TGFβ and IL10 (6). Tfh cells promote B cell activation and differentiation, stimulate generation of long-lived antibodies, express BCL6, IRF4, MAF, and BATF, and secrete IL21, IL4, and IL10(7).

T-cell responses are essential to health maintenance but also contribute to pathogenesis. Reduced Treg cell numbers or functions are seen in many autoimmune diseases as exemplified by association of Foxp3 gene mutation in some patients with IPEX (immune dysregulation, polyendocrinopathy, enteropathy, X-linked) syndrome (1). Indeed, balancing the functions of effector and regulatory T-cells appears critical for promoting favorable outcomes. Auto-reactive T-cells that escape negative selection in the thymus are involved in the development of autoimmune diseases such as type 1 diabetes (T1D) (8). Polymorphisms at HLA-DR and –DQ class II loci strongly associate with increased risk for T1D (9), which is notable since these genes encode proteins involved in presentation of antigenic peptides to T-cells. Thus, tools and approaches for characterizing T-cells and monitoring their function over time are paramount to the development of preventative therapeutic strategies for autoimmune diseases. They are also necessary for evaluating responses elicited by prophylactic vaccines as well as a rapidly expanding array of immune modifying agents used in the treatment of chronic conditions such as arthritis and more recently cancer (10).

Traditionally T-cells have been characterized using a limited number of cell-surface markers, transcription factors, and secreted cytokines, which are measured using flow cytometry and other antibody-based assays (11). Classification of T-cell responses using such a knowledge-based approach is inherently biased. Systems approaches could instead yield an unbiased molecular classification since they rely on the use of high-throughput profiling technologies to measure constituents in a given biological system. Recent technological advances allow, for instance, the genome-wide profiling of transcript abundance at the population and single-cell levels, the assessment of immunodominant antigens by pathogen proteome microarrays, and the determination of antibody and TCR repertoire diversities by DNA sequencing (12).

Transcriptome profiling has been leveraged successfully to investigate pathogenesis (13-15), innate immunity (16), and responses to vaccines (17, 18). Systems approaches in human immunology studies have largely consisted in profiling abundance of cellular RNA in whole blood or peripheral blood mononuclear cells (PBMCs) of the study subjects. Only seldom have whole transcriptome readouts been used in *in vitro* transcriptional assays (19-21).

The work presented here employs transcript profiling for the unbiased characterization of T-cell responses. As a proof of principle, we used microarray datasets to characterize transcriptome-wide responses to immunodominant peptides in over 300 PBMC cultures. This meta-analysis identified co-expressed gene clusters that categorize antigenic responses using a purely data-driven approach. While the limited diversity of the antigenic repertoire tested and lack of immune phenotyping data constrains the interpretation of the findings described in this article, the strategy that we present can serve as a basis for further studies that will establish unbiased classification of antigen-specific responses. These may contribute to further expand our understanding of T-cell immunobiology and serve as a foundation for the development of a new generation of immune profiling assays.

## Materials and Methods

### Cell culture and peptide stimulation

Blood samples were derived from subjects participating in studies under the auspices of Control and Diabetes Registries and Infectious disease registry. Informed consent was obtained from all subjects according to IRB-approval protocols at Benaroya Research Institute (Seattle WA) and at Baylor Research Institute (Dallas, TX). PBMCs were isolated by ficoll method (GE Healthcare Bio-Sciences Corp., Piscataway, NJ), and were frozen until use. Immunodominant peptides used in this experiment consisted of allergy peptides: *Alnus glutinosa* major allergen Aln g 1 derived peptide Alng1p10 and *Felis domesticus* allergen Fel d 4 derived peptide Feld4p30, microbial peptides: Candidap38 from *Candida albicans*, CMVp86 from cytomegalovirus, BHp28, BHp46, H1p21, H1p33, H1p34, H1p45, H3p13, H3p3, H3p306, H5p86, MPp54, MPp8, MPp6, NPp125, NPp1528, NPp68, and Flu-MP from influenza virus, and WNVEp7 and WNVNS2ap2 from West Nile virus, and type 1 diabetogenic peptides: GAD65C1, GAD65p73, IGRPC4, IGRPp11, IGRPp12, IGRPp13, IGRPp14, PPIC3, ZnT8C5, ZnT8p15, ZnT8p16, ZnT8p17, ZnT8p18, ZnT8p33, ZnT8p65, ZnT8p68, and ZnT8p93. Peptide sequences are provided in **S1 Table**. Peptides were purchased from Mimotopes (Victoria, Australia), and were dissolved in 100% DMSO or 50% acetonitrile. PBMCs were thawed, washed, and cultured in 14 ml polypropylene tubes (BD Falcon, Tewksbury MA) with either RPMI 1640 (GIBCO, Grand Island NY) supplemented with 10X AB serum (Gemini Bio-Products, West Sacramento CA), 50 uM of 2-mercaptoethanol (Sigma, St. Luis MO), 25 ml of HEPES (GIBCO, Grand Island NY), 1% sodium pyruvate (Sigma, St. Luis MO), 1% non-essential amino acid (Sigma) or RPMI 1640 supplemented with 1X serum replacement (Sigma, St. Luis MO), 1 mM sodium pyruvate (Sigma, St. Luis MO), 1x penicillin and streptomycin (Sigma, St. Luis MO). PBMCs were stimulated with peptides for 24 hours. The same amount of acetonitrile or DMSO without peptide was added to medium in control cultures (medium control). Subsequently cells were washed with PBS and lysed in RLT buffer (Qiagen, Valencia CA) supplemented with 1% 2-mercaptoethanol (Sigma, St. Louis MO).

### Microarray

RNAs were extracted using the RNeasy Mini Kit (Qiagen, Valencia CA), were quantitated by NanoQuant (Tecan, Männedorf, Switzerland), and their qualities were assessed on a Bioanalyzer 2100 (Agilent, Santa Clara CA). Samples were then labeled and amplified using Illumina TotalPrep-96 RNA amplification kit (Ambion, Grand Island NY). Finally, samples were hybridized to Illumina HumanHT-12 v3 or v4 bead chips, and read on an Illumina HiScanSQ scanner (Illumina, San Diego CA). Background subtracted data were generated using GenomeStudio (Illumina, San Diego CA).

### Microarray data analysis

Data were analyzed in R (22). The data were preprocessed by quantile normalization, and flooring values <10 to 10. Common probes between the two microarray chip versions were selected. Probes present in less than 10% of all the samples (PALX10%), and probes with difference between minimal and maximal expression across samples less than 100 (range 100) were excluded.

The ratio and difference between stimulated samples and medium controls from the same donor were computed. Experiment 5 consisted of cultures in triplicates for each peptide stimulation and medium control, and the mean of expression from the triplicate culture was used for the ratio and difference calculations. Filtered data consisting of probes with at least 5 samples and samples with at least 50 probes, with absolute log_2_(ratio) and absolute differences greater than 1 and 200, respectively, were selected for clustering. To cluster probes and samples, ratio values less than 0.667 were set to-1, values between 0.667 and 1.5 were set to 0, and finally values equal or greater than 1.5 were set to 1. Probes and samples were individually clustered using Hartigan’s *k*-Means (23) for all *k* in the range of 1-100, inclusive, and in the range of 1-30, inclusive, respectively. The Jump algorithm (24) was used to determine the final k = 9 and k = 11 for probes and samples, respectively. Resulting clusters were called gene clusters and sample sets for probes and samples, respectively. The lists of genes related to immune functions were obtained from ImmPort (http://immport.niaid.nih.gov) (data retrieved in January 2012), the *HUGO Gene Nomenclature Committee at the European Bioinformatics Institute* (http://www.genenames.org/genefamilies/CD) (25) (data retrieved in March 2012), and MetaCore (GeneGO Inc, MI, [www.genego.com]) (data retrieved in January 2012). The biosets were created by matching the Illumina probes with the gene symbols from the lists of genes (**S2 Table**). Thus, the biosets were not derived from our data. The transcription factor (TF) bioset included genes that encode for not only TFs, but also regulators of transcription.

Enrichment of genes in gene clusters that were also in biosets, and enrichment of peptide types in sample clusters were determined by Fisher’s exact test at a significance level of 0.05.

For a given gene cluster and a sample set, the mean of ratios for the probes was taken for each sample. Then, the mean of the means of ratios was taken, transformed into log_2_ scale, and assessed the mean of the means of ratios was greater than 1 or less than-1 by one-sample test (one-sided) at a significance level of 0.025 to examine if the given sample set showed significant change in expression for the given probes.

### PubMed literature search

TFs found in both TF bioset and our filtered data were identified, and they were individually searched in PubMed (http://www.ncbi.nlm.nih.gov/pubmed), using both gene symbols and synonyms, for associations with helper and regulatory T cells, including Th1, Th2, Th9, Th17, Th22, Treg, and Tfh cells. For the CAT gene, many false positives were returned using the symbol CAT, so the official full name Catalase was used instead of the symbol CAT. All searches were limited within the title and abstract using the [TIAB] tag. To narrow the search results to journal articles in humans, PubMed sidebar filters “Journal Article” and “Humans” were activated. Some synonyms were excluded from the search because they were not specific to the genes. For example, “DELTA” a synonym for YY1 and “MAIL” a synonym for NFKBIZ were excluded. The number of articles returned from the search was recorded to assess the extent to which TFs found in our filtered data were studied in the context of the various T-cell types. In addition, if an article was associated with more than one T-cell types, it was also recorded in “Overlap”. Moreover, a control set of 20 TFs (BANP, BNIP3, CTBP1, ELF5, FOSL2, GLIS2, HAND1, LOC642559, MOS, OVOL1, PBX2, POU5F1, PRDM16, RARA, RBMS1, SATB2, STAT5A, STOX1, TTF1, ZBTB38) were selected from the TF bioset randomly based on arbitrary integers chosen by using a function sample() in R (22). PubMed searches were performed for these TFs as described above. Data were retrieved in March 2014.

### Functional annotation by DAVID

Each gene cluster was annotated using DAVID (data retrieved in October 2013 using DAVID v6.7, http://david.abcc.ncifcrf.gov/home.jsp) (26, 27). The common probes between Illumina HT12 v3 and v4 (39426 probes) were used as a background gene list for enrichment analysis for Gene Ontology biological process (GO BP). Statistical significance of associated BPs was assessed using the FDR adjusted p-value at a significance level of 0.05.

### Results

#### Measuring *in vitro* transcriptome responses to antigenic peptides

Traditionally T-cell responses have been characterized and classified using limited panels of molecular markers selected based on *a priori* knowledge about T-cell immunobiology. Our goal was to identify and characterize antigenic-peptide specific T-cell responses in an unbiased manner using a data-driven approach. Hence, we set out to utilize transcriptome profiling to capture the breadth of the response to antigenic peptides. Changes in transcript abundance were measured on a genome-wide scale in a set of 352 peptide-stimulated samples. Cryopreserved PBMCs from various donors were stimulated *in vitro* with a broad range of antigen-derived peptides for 24 hours. Changes in RNA abundance were measured using Illumina Beadarrays. A total of 7 independent experiments were included in the analysis: Experiments #1-3 examined samples for which cytokine responses measured by a multiplex protein assay were known. In these experiments PBMCs from 1∼3 donors were stimulated with peptides derived from microbes, alder and cat allergens, and type 1 diabetogenic (T1D) proteins. Experiment 4 compared influenza virus (flu) peptide-induced responses in young and old individuals. This study involved 11 donors and 13 flu derived peptides. Experiment 5 examined PBMCs from a single donor, exposed to flu derived peptides and simultaneously measured the effect of freezing PBMCs on the peptide specific response at different T-cell precursor frequencies. Finally, experiments 6 and 7 examined differences in T1D or flu matrix protein derived peptides-induced responses in individuals with T1D and age/gender/HLA-matched healthy controls (28). These two studies involved 35 patients and 30 controls in total. The T1D peptides were derived from the 65kDa isoform of glutamic acid decarboxylase (GAD65), islet-specific glucose-6-phosphatase catalytic subunit-related protein (IGRP), preproinsulin (PPI), and zinc transporter 8 (ZnT8). The antigen sources of peptides used in those experiments are listed in **Table 1**.

**Table 1.** List of peptides used in PBMC cultures.

| Disease | Source protein | Peptide |  |  |  |  |  |  |
| --- | --- | --- | --- | --- | --- | --- | --- | --- |
|  |  | Exp1 | Exp2 | Exp3 | Exp4 | Exp5 | Exp6 | Exp7 |
| Allergy | *Alnus glutinosa* major allergen Aln g1 |  |  | Alnglp10 |  |  |  |  |
|  | *Felis domesticus* allergen Fel d 4 |  |  | Feld4p30 |  |  |  |  |
| T1D | GAD65 |  | p73 |  |  |  | C1 |  |
|  | IGRP |  |  |  |  |  | C4, p11, p12, p13, p14 | C4, p11, p12, p13, p14 |
|  | PPI |  |  |  |  |  | C3 | C3 |
|  | ZnT8 |  | p68, p33, p65, p93 |  |  |  | C5, p15, p16, p17, p18 |  |
| Microbial | *Candida albicans* | p38 |  |  |  |  |  |  |
|  | Cytomegalovirus (CMV) | p86 |  |  |  |  |  |  |
|  | Influenza virus (Flu) | H1p45,<br>H5p86 |  |  | BHp28, BHp46,<br>H1p21, H1p33,<br>H1p34, H3p13,<br>(H3p3, H3p306),<br>MPp6,<br>(MPp8, MPp54),<br>NPp125, NPp68 | MPp8,<br>MPp54,<br>NPp1528 | MP |  |
|  | West Nile virus (WNV) |  | Ep7,<br>NS2ap2 |  |  |  |  |  |

Peptides that were used to examine antigenic responses in PBMC cultures are listed in this table. GAD = glutamic acid decarboxylase, IGRP = islet-specific glucose-6-phosphatase catalytic subunit-related protein, PPI = preproinsulin, ZnT8 = zinc transporter 8. C1, C3, C4, and C5 were pools of peptides derived from GAD65, IGRP, PPI, and ZnT8, respectively. Influenza A HA, and influenza B HA proteins are denoted as H, and BH, respectively. Peptides (H3p3, H3p306) and (MPp8, MPp54) were pools of 2 peptides H3p3 and H3p306, and MPp8 and MPp54, respectively. MP was pool of overlapping peptides encompassing the entire influenza A M1.

#### Identification of peptide responsive gene clusters

We first verified that the data obtained from the different experiments could be consolidated in a single meta-analysis. Principal component analysis (PCA) was performed using 12069 probes that showed variability based on filter criteria for selection of transcripts detected in at least 10% of all samples (PALX10%) and with range 100 (see Methods) across 352 stimulated samples and 83 medium controls. As could be expected, the resulting plot showed a clear separation of samples between independent experiments (**Figure 1A**). However, such a separation was not present once stimulated conditions were normalized to their respective medium controls (**Figure 1B**). This finding demonstrates that the use of respective medium controls as a common denominator across the different experiments is an effective approach for normalizing our data and controlling batch effects. This normalization method also made it possible to analyze changes induced by stimulation regardless of baseline variations among donors. Thus, this strategy of using the normalized data to each donor’s baseline was adopted for carrying out the meta-analysis presented in this article.

**Figure 1.**
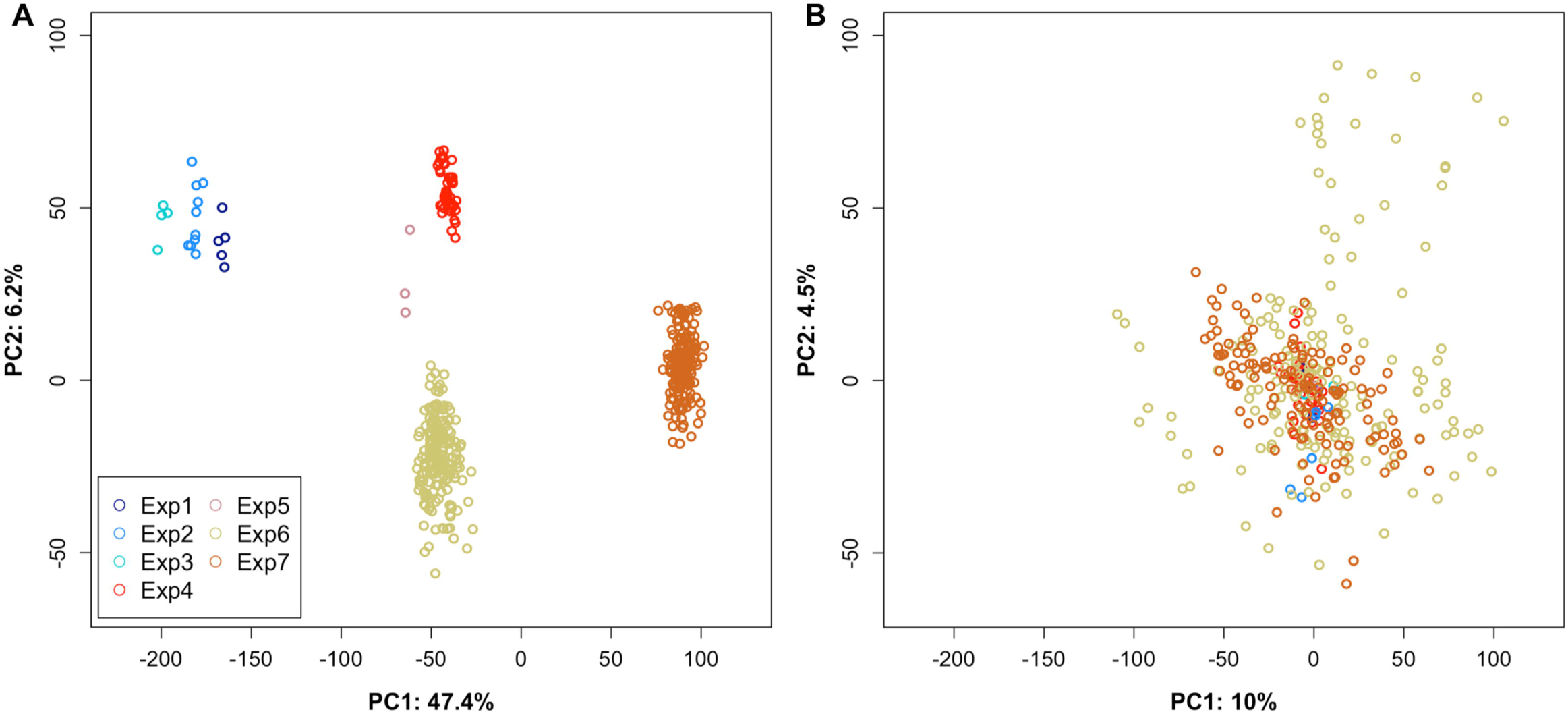
Principal component analysis of transcriptional profiles before and after normalization to medium controls. 12069 probes, after exclusion of probes with little variations across datasets, were used as input in the PCA. Samples were color-coded by experiments. Percent variations explained by PC1 and PC2 are shown. A) Before data normalization: PCA of data from 352 stimulated samples and 83 medium controls. Intensity values in log_2_ scale were used. B) After data normalization: PCA of data from 352 samples. Log_2_(ratio) data after normalization of stimulated samples to their respective medium controls were used.

We sought to categorize T-cell responses in an unbiased fashion, based on transcriptional patterns observed following antigenic peptide stimulation. For this we employed an unsupervised clustering approach grouping samples and genes according to patterns of responsiveness, which is described in detail in the Methods section. Clustering methods are known to be prone to noise. Thus, following normalization of stimulated conditions to their respective medium controls, both probes and samples were filtered on the same cutoffs; ratio of 2 for upregulation or 0.5 for downregulation and absolute difference of 200. The probes were retained if at least 5 samples passed those cutoffs, and the samples were retained if at least 50 probes passed those cutoffs. The filtered data consisted of 949 probes and 111 samples while the data prior to the filtering consisted of 12069 probes and 352 samples. To identify co-expressed genes, this dataset was clustered two-ways (i.e. probes and samples) using Hartigan’s K-means and applying Jump theory as published previously (24). The clustering resulted in 9 gene clusters (C0-C8) and 11 sets of samples (SS0-SS10) (**Figure 2**). The complete cluster information for both probes and samples can be found in **S3 and S4 Tables**, respectively.

**Figure 2.**
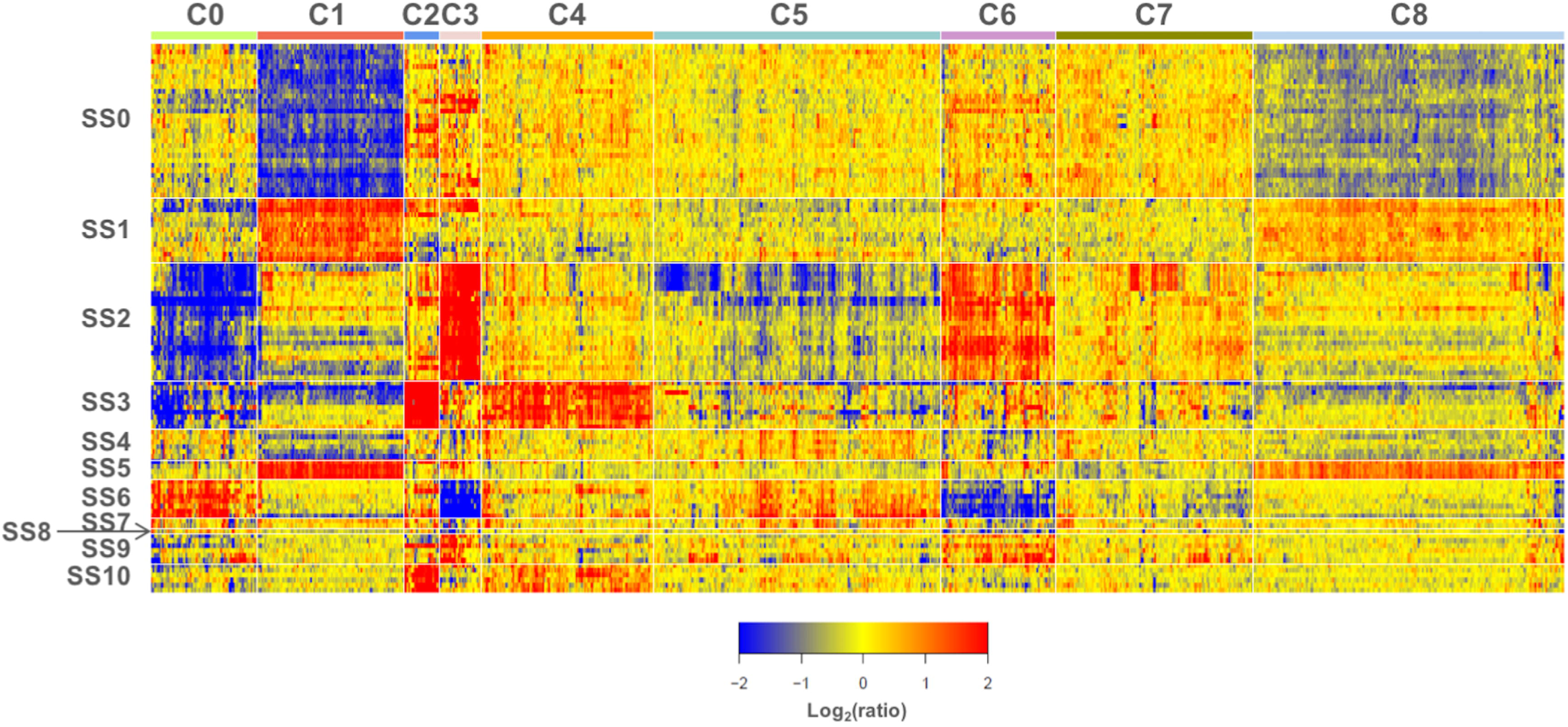
Heatmap of co-expressed genes. Log_2_(ratio) (stim/non-stim) of 949 probes and 111 samples that passed the filter were visualized on this heatmap. Probes and samples were clustered by K-means clustering using the Jump method. Probes were arranged in columns and samples in rows. Vertical and horizontal white lines divide probes and samples in clusters, respectively. Red, blue, and yellow indicate an increase, decrease, and no change, respectively, in abundance over the medium controls.

#### Functional enrichment analysis of peptide-responsive gene clusters

Next, the gene clusters defined above were characterized functionally. Although we are attributing changes in transcript abundance to regulation in T-cells, these changes could also be caused by other cell types that are exposed to T-cell factors in PBMC culture.

Gene clusters were functionally characterized at a high level through Gene Ontology (GO) enrichment analysis using the DAVID annotation tool (26, 27). The top five significant biological process terms associated with each cluster at a significance level of 0.05 using the FDR adjusted p-values are summarized in **Table 2**. Cluster 0 (C0) was associated with lipid metabolic process and response to external stimulus. C1 was associated with translation and metabolic process. C2 through C7 were associated with various aspects of immune response. No GO terms were significantly associated with C8.

**Table 2.**
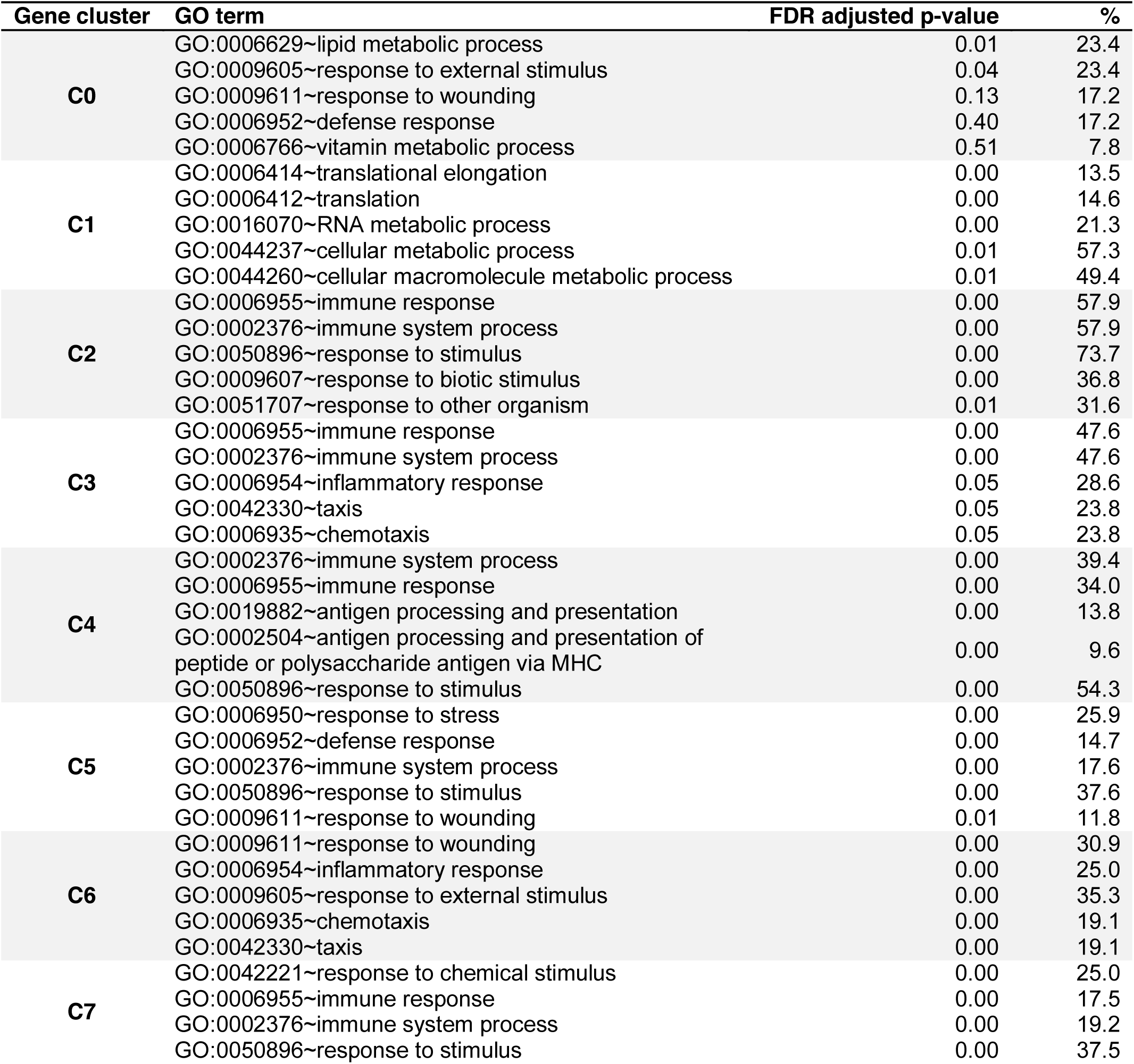
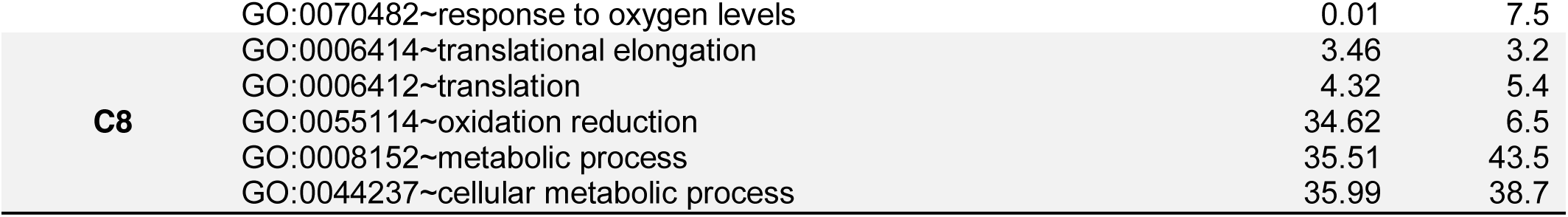
Top five GO biological process terms associated with gene clusters.

DAVID functional annotation was performed using lists consisting of probes from each cluster and background of 39426 probes, which were considered for ratio and difference filtering. GO biological process terms associated with each cluster were ordered by p-values and the top 5 GO terms were listed in this table. The FDR adjusted p-values and % were rounded to 2 and 1 decimal points, respectively. % indicates percentage of genes associated with a term / total # of query genes. Only 2 terms were significant for C0, 3 terms for C3, and none for C8.

Next we refined our interpretation by mapping co-clustered probes to several relevant functional categories (biosets) corresponding to transcription factors (TF), Cluster of Differentiation (CD) molecules, cytokines and chemokines, cytokine and chemokine receptors, T-cell and B cell receptor (TCR and BCR) signaling, and antigen processing and presentation (**Figure 3**). These functional gene lists were formed based on information compiled from the GeneGO MetaCore knowledgebase (GeneGO Inc, MI, [www.genego.com]), HGNC Database (http://www.genenames.org/genefamilies/CD) (25) and Immport (http://immport.niaid.nih.gov), and were summarized in **S2 Table**. Overlaps existed between these biosets due to the nature of the annotation of the genes. For example, CSF2, IFNG, IL2, IL4, IL5, IL10, and TNF were all categorized as cytokines, but were also in TCR signaling; and were therefore found in both the cytokine and TCR signaling biosets. Immune bioset-derived probes accounted for greater than 30% of the probes constituting each cluster with the notable exception of C1 and C8 (≤10%). Over 60% of the probes constituting C3 were found in the immune biosets. Using the Fisher’s exact test (H_a_: odds ratio > 1 at a significance level of 0.025), we determined that C0 was significantly enriched for CD molecules, C3 for cytokines, C4 for molecules found in the antigen processing, C5 for CD molecules, C6 for cytokines. In addition, presence of TFs, cytokines, and chemokines in each cluster is summarized in **Table 3**.

**Figure 3.**
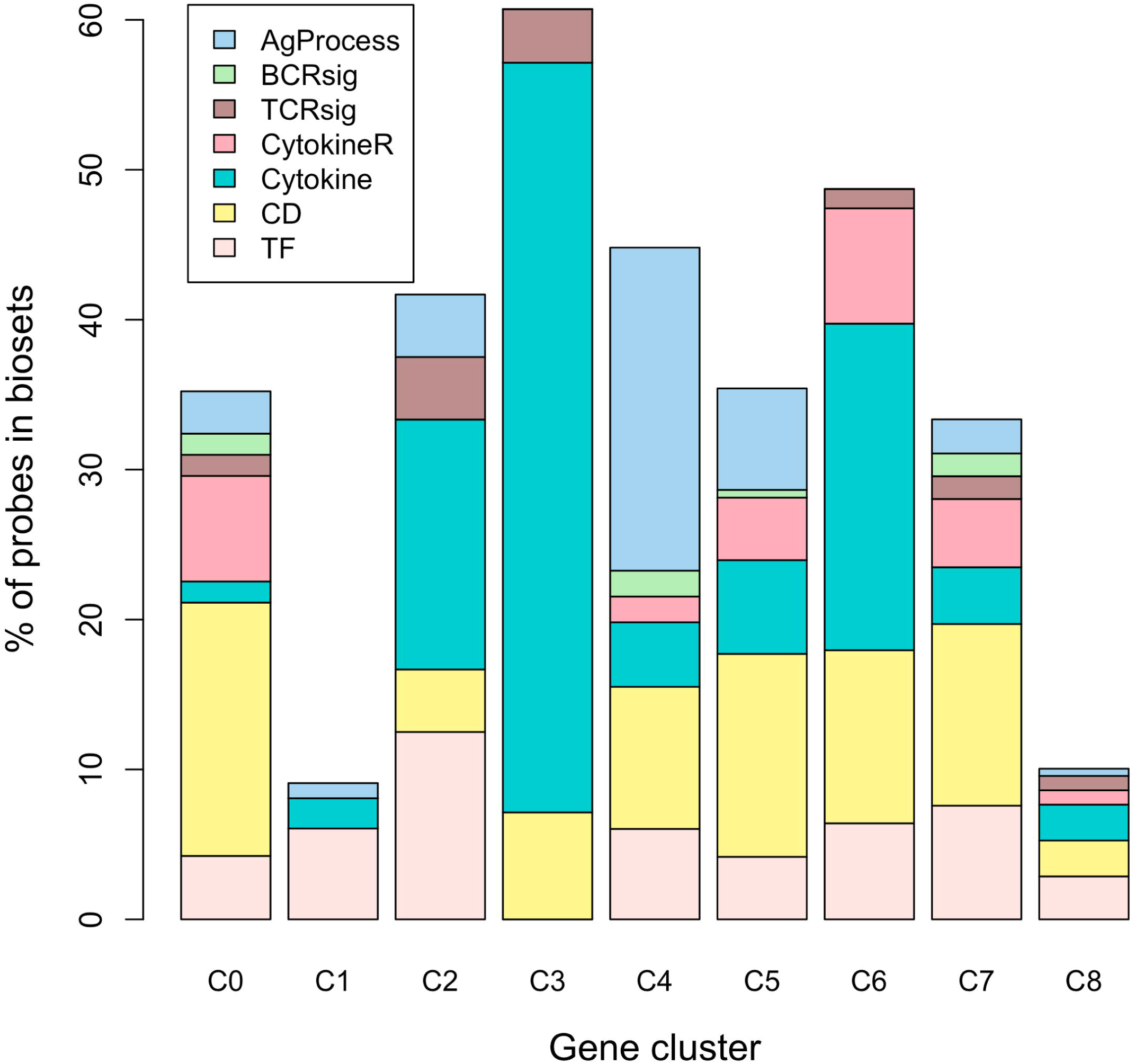
Immunological annotation of gene clusters. TFs, CD molecules (CD), Cytokine, cytokine receptors (CytokineR), BCR signaling (BCRsig), TCR signaling (TCRsig), antigen processing and presentation (AgProcess) biosets were created by matching the gene symbols obtained from GeneGo, HGNC, and Immport websites with Illumina probes. Gene clusters are shown along the x-axis. Percent of genes in each cluster that overlap with the biosets are shown along the y-axis.

**Table 3.**
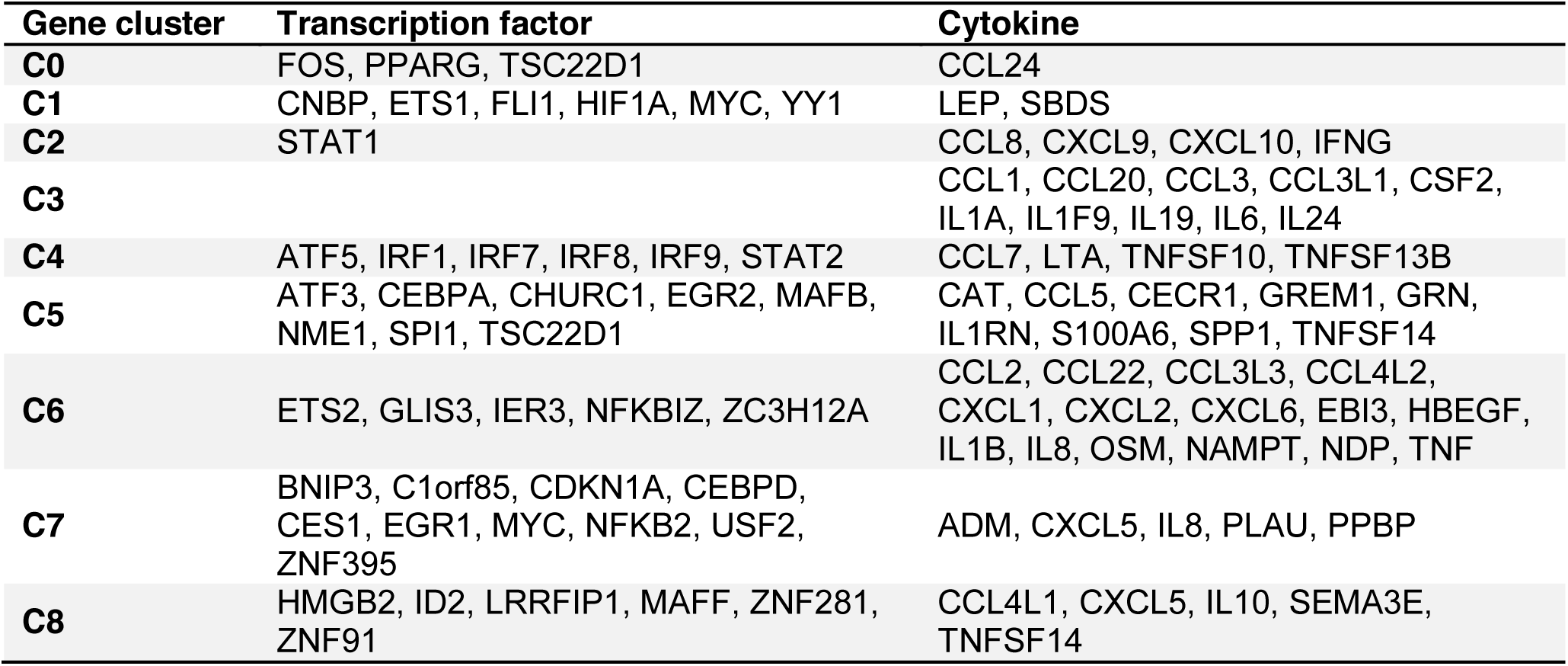
TFs and cytokines found among gene clusters C0 to C8.

This table provides the gene symbols of transcription factors and cytokines in gene clusters responsive to peptide stimulation in PBMC cultures. Some genes were in multiple clusters because multiple Illumina probes could be associated with the same gene. When multiple probes targeting the same gene were found within a cluster, the gene symbol was listed only once.

#### Assessing knowledge gaps among peptide-responsive gene clusters

We identified 43 transcription factors, regulated in response to peptide stimulation in our assay system, and assessed whether these were already known to be associated with T-cell function. Literature searches were conducted in PubMed to determine the frequency of articles indexed for these transcription factors that contained in their titles or abstracts keywords describing CD4^+^ T-cell phenotypes, “Th1”, “Th2”, “Th9”, “Th17”, “Th22”, “Treg”, and “Tfh”. The results are shown in **Figure 4**. Out of 43 transcription factors, 9 (21%) had at least 5 unique articles associated with T-cell phenotypes, 60% had at least one. When PubMed searches were carried out in a similar manner for 20 transcription factors that did not belong to any of our gene clusters and were selected at random, only 2 of them (10%) returned more than 5 articles containing those keywords. The fact that the signatures that we observe were enriched in transcription factors known to be relevant to differentiation and function of T-cell was not altogether surprising. However, the wide spread in the number of T-cell types associated-articles returned across all 43 genes indicate that the degree to which those genes have been investigated in the context of T-cell immunobiology varied greatly. While searches for STAT1 returned over 200 articles associated with different types of T-cell responses, transcription factors such as PPARG and ATF5 returned fewer than 5 articles related to T-cell phenotypes, and 14 transcription factors returned none. Those transcription factors that displayed limited or non-existent overlap with literature on CD4^+^ T-cell types should be considered for further investigation as potential candidate master regulators of T-cell differentiation and effector functions (according to the principle of “guilt by association”). One of those candidates was ATF5, which belongs to cluster C4, annotated as immune system process, immune response, and antigen processing and presentation (**Table 2**). ATF5 is a member of cAMP response element binding (CREB)/activating transcription factor family, and regulates cell differentiation, proliferation and survival (29). Only one journal article found by our PubMed search indicates an association of ATF5 with Th1 response. In the article, Chuang *et al*. showed ATF5 was induced by phytohemagglutinine and a combination of anti-CD3 and anti-CD28 antibodies in T-cells, and that the induced ATF5 expression was associated with increased IFNG and TNFA expressions (30). Another article related to ATF5 but not in association with T-cell response suggests that ATF5 induces expression of mammalian target of rapamycin (mTOR) through PI3K/AKT pathway in BCR/ABL-transformed 32D cells (31). PIK3/AKT pathway can signal through mTOR, which in turn participate in regulation of T-cell differentiation and proliferation (32). Therefore, it is conceivable that ATF5 may participate in the establishment of an antigen-specific T-cell response by increasing Th1 cytokine secretion as well as by activating mTOR to control T-cell fate through PI3K/AKT pathway downstream of TCR activation. Altogether our analysis provides the potential for discovery of novel factors involved in T-cell differentiation and effector function.

**Figure 4.**
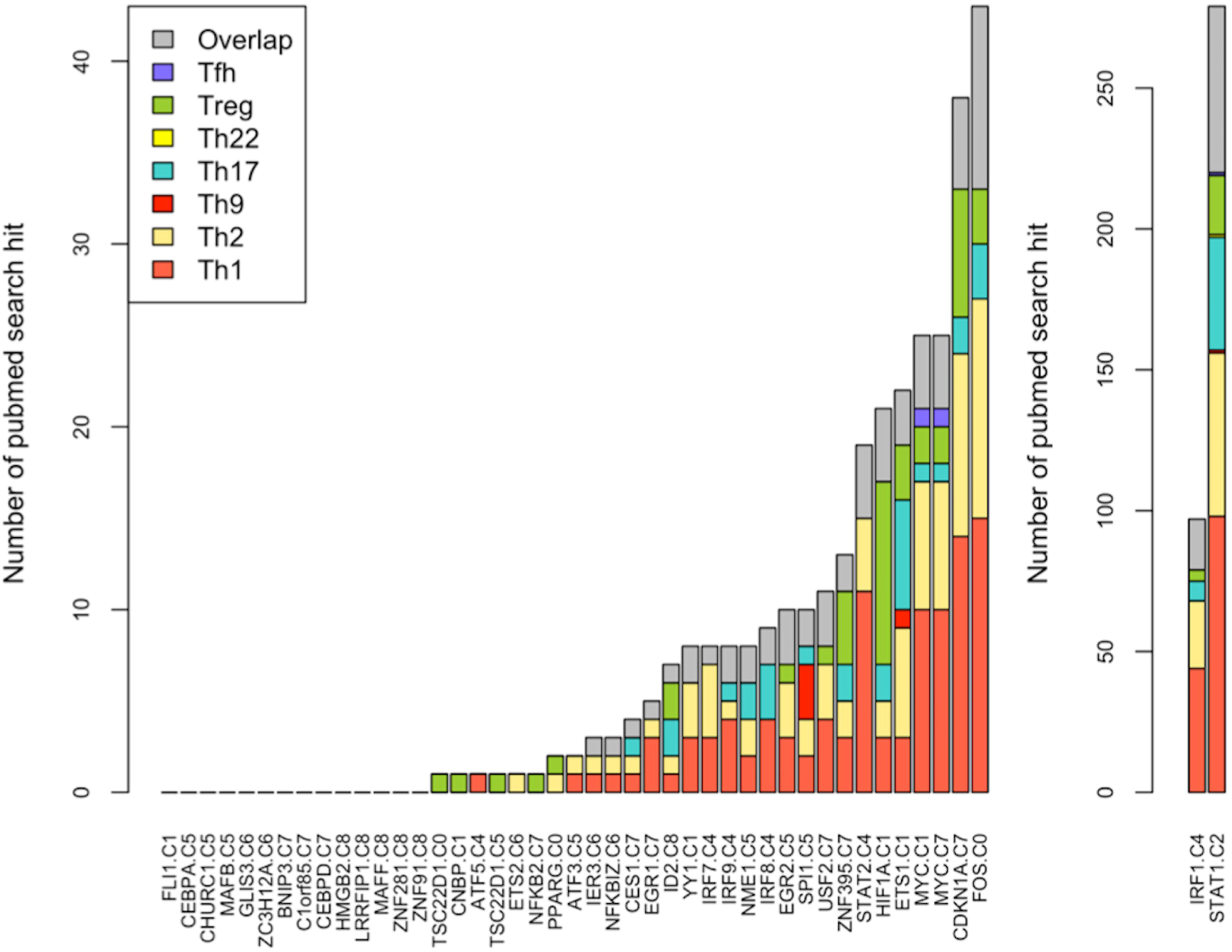
Literature profiles of TF genes. T-cell literature profiles for TF genes were generated. Gene symbols and gene cluster information are shown along the x-axis. The numbers of returned articles were plotted. If an article was associated with more than one T-cell types, it was also counted in “Overlap.”

#### A novel unbiased definition of types of T-cell responses

Our study aimed at leveraging systems approaches to arrive at an unbiased (i.e. data-driven) classification of T-cell responses obtained after peptide stimulation in an *in vitro* system. Unsupervised filtering and clustering identified a collection of gene clusters that we have characterized functionally. To classify the observed responses, one-sample t-tests were performed for each gene cluster (C1 to C8) and sample set (SS1 to SS10) combinations, except for SS7 and SS8 that consisted of fewer than 3 samples. The change in mean expression was considered significant if mean expression change was > 2 or < 0.5 at a significance level of 0.025 (one-sided). The results, summarized in **Table 4** and shown in **Figure 2**, indicate that each sample sets exhibited expression of one or a combination of two or more gene clusters, except SS4 that did not show significant change in any gene cluster. We designated downregulation and upregulation of gene clusters with superscripts “lo” and “hi”, respectively, following the gene cluster names. For instance, the SS2 response was defined by changes in 3 different gene clusters and was noted as C0^lo^C3^hi^C6^hi^. Overall 8 different types of T cell responses were identified in this manner (**Table 4**). On the basis of this classification we then compared responses to type 1 diabetogenic peptides (88 samples) and microbial peptides (21 samples) (**Figure 5**). For each category of the peptides, the percentages of the samples among the total number of samples in a given peptide category was computed for each sample set. Then, the sample set was labeled by the responses defined in Table 4. We found the responses elicited to be distinct between these two categories of peptide stimulated samples. Type 1 diabetogenic peptide induced responses were dominated by the types C1^lo^ (35% of the samples), C0^lo^C3^hi^C6^hi^ (24%) and C1^hi^ (15%), while microbial peptide stimulation elicited C2^hi^C4^hi^ (33%), C3^hi^ (19%), C2^hi^ (14%), C0^hi^C3^lo^ (14%), and C0^lo^C3^hi^C6^hi^ (14%) patterns. Notably, only the later type, C0^lo^C3^hi^C6^hi^, was found represented at a high proportion in both stimulation groups. Functional interpretations for the different types of T cell responses identified via this approach are provided below.

**Figure 5.**
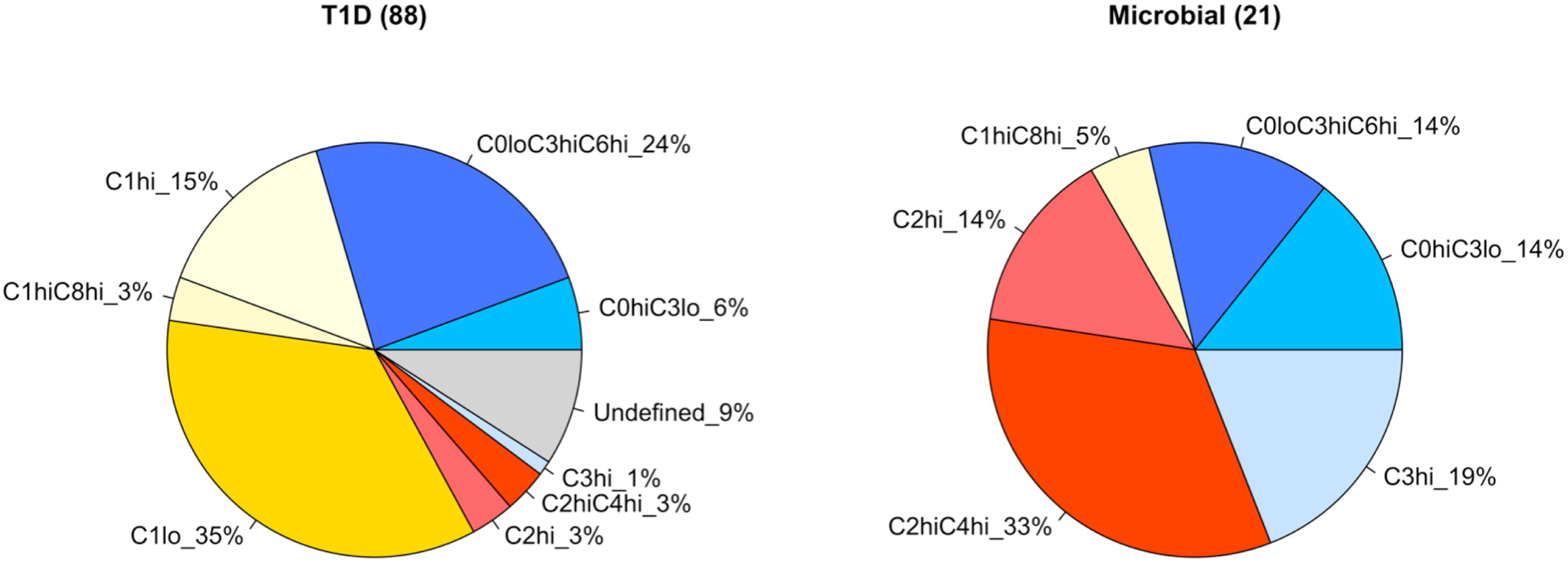
The distribution of T-cell responses for T1D and microbial peptides. Peptides were categorized into Type 1 diabetogenic and microbial proteins: labeled in the figure as “T1D” and “Microbial”, respectively. Percentage of samples that exhibited T-cell response types determined in Table 4 were computed and shown in the pie chart. Each pie represents a T-cell response type and its percentage. Numbers of samples for each peptide category are shown in parenthesis on the top of each pie chart. Undefined includes SS7 and SS4.

**Table 4.**
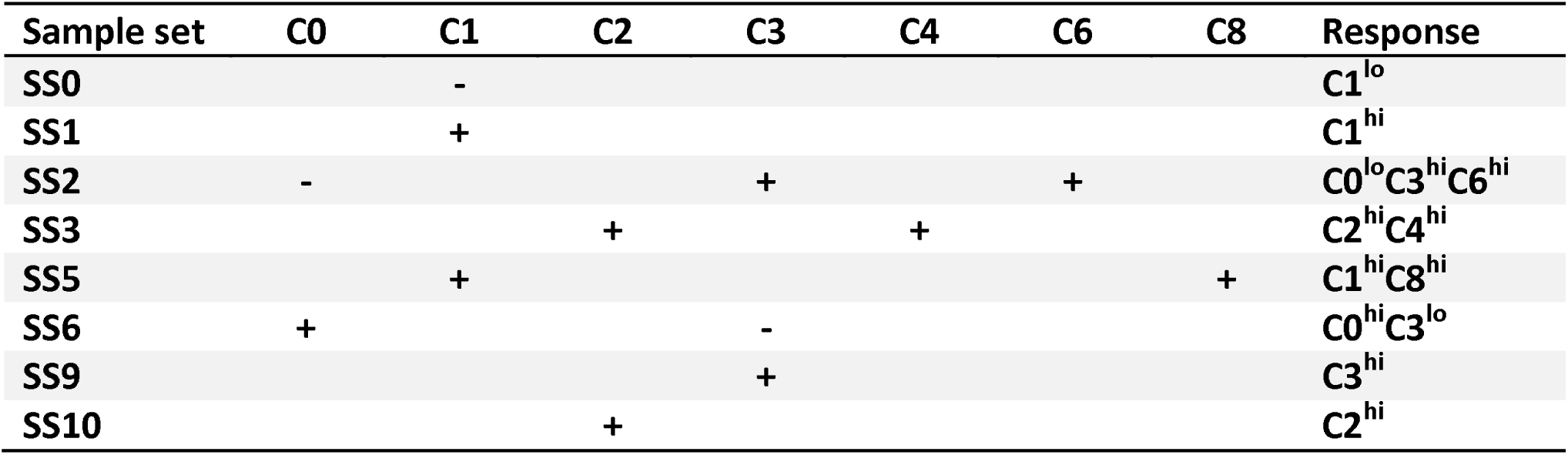
Combination of gene clusters used for characterizing the types of responses observed.

+ and-in the table indicate that mean expression change of a sample set for a gene cluster was > 2 and < 0.5, respectively, at a significance level 0.025 in one sample t-test (one-sided). Blank cell indicates that the mean expression change of a sample set for a gene cluster was neither > 2 nor < 0.5 at the significance level. SS7 and SS8 were not tested as they consisted of only two and one samples. SS4, C5, and C7 were omitted from the table for lack of significant change in expression.

#### Functional characteristics of T1D peptide responses

Cluster C1 signatures dominated the responses to T1D peptides, with C1^lo^ and C1^hi^ responses representing 35% and 15% of the response types, respectively. The C1 cluster consisted of 99 probes and was enriched with genes involved in translation and RNA metabolic process. Those genes included ribosomal proteins RPL7, RPL9, RPL23, RPLP1, RPS28 for translation, and CROP, ETS1, HIF1A, HNRPC, HSPA1A, LCOR, MYC, NOP56, PABPC1, PABPC3, RPL7, RPS28, SBDS, SFRS11, SLBP, ZFP36L1 for RNA metabolic processes such as pre-mRNA processing, mRNA stability and translation. Previously published studies suggested that translational regulation is important for control of inflammation. For example, ZFP36 destabilizes TNFα mRNA after macrophage activation (33); a ribosomal protein RPL13A in murine macrophage functions in resolution of inflammation (34); and translation of inhibitors of NFκB and post-transcriptional suppressors of cytokine expression are increased at early cell activation upon LPS stimulation of murine macrophage (35). Although the specific genes described in these previous studies are different from those identified in this study, it is possible that C1 genes are also involved in the translational regulation of the inflammatory response. The C0^lo^C3^hi^C6^hi^ type was the other predominant response observed, with 24% of the T1D peptide responses being of this type. This type of response was characterized by a concomitant suppression of an anti-inflammatory program (C0) and induction of pro-inflammatory ones (C3 and C6). C0 consisted of 71 probes, and included the genes APOC1, APOE, CD36, and PPARG. PPARG, prevents T-cell activation via reduction in IL2 production, and inhibits macrophage activation and cytokine production by monocytes (36), while APOE suppresses T lymphocytes proliferation (37). Notably C0 also contained CD163, CSF3R, and LYZ genes. CD163 is an anti-inflammatory monocyte/macrophage marker (38). CSF3R activates STAT5 that targets SOCS3, which in turn modulates IFNG response (39). LYZ encodes lysozyme that is known for its antimicrobial activities. Overall, this transcriptional program may be associated with suppression of immune responses. Conversely C3 can be associated with pro-inflammatory function. This cluster consisted of 28 probes, and the genes in C3 associated with inflammation were mostly cytokine genes, which were also observed in the biosets-analysis. These genes included CCL1, CCL20, CCL3, CCL3L1, CSF2, IL1A, IL1F9, IL6, and IL19. Activated T-cells secret large amounts of CCL1, while CCL1 receptor, CCR8, is expressed on subsets of T-cells including Th1, Th2, and Treg cells (40). CCL3 and CCL3L1 are chemoattractants that promote the inflammatory response by recruiting immune cells of the myeloid lineage (41, 42). CCL20 expression is induced by proinflammatory cytokines, and recruits immature DCs and macrophages to inflammation sites through interaction with CCR6 (43). CSF2 is a pro-inflammatory cytokine, which supports survival of Th17 cells (44). The pro-inflammatory cytokine IL6 induces acute phase proteins, and regulates recruitment of immune cells to inflammation sites. It also regulates the balance between Th17 and Treg cell differentiation, and indirectly antibody production by B cells (45). IL19 increases Th2 cytokine production, and induces expression of IL6, IL8 and IL10 in monocytes (46). Overall, C3 genes may promote inflammation. C6 was associated with the GO term “inflammatory response.” This cluster consisted of 78 probes, and the genes in C6 enriched for this term were mostly cytokine and cytokine receptor genes, which were also observed with the biosets-analysis. Some of these genes were pro-inflammatory cytokines (e.g.: IL1B and IL8). Others such as CCL2 and TNF can be both pro- and anti-inflammatory. CCL2 recruits monocytes, memory T-cells, and NK cells, but also can negatively regulate macrophage activation and pro-inflammatory cytokine production, and skew macrophage polarization towards M2 with anti-inflammatory activity (47). TNF activates caspases, NFκB and MAPK pathways for induction of inflammation, but also can increase anti-inflammatory factors such as IL10 and corticosteroid to downregulate its expression (48, 49). The cytokine receptors in C6 include FPR2 and TNFRSF4. Activation of G-couple protein receptor FPR2 may result in pro-or anti-inflammatory response. The ligands for this receptor include anti-inflammatory lipids lipoxin A_4_, and serum amyloid A, and their bindings to the receptor induce apoptosis and survival signal, respectively (50). TNFRSF4 is a TNF receptor family member, which binds to OX40L expressed on APC. It is preferentially expressed on activated regulatory CD4^+^T-cells, NKT-cells, NK cells, and neutrophils. The signaling through TNFRSF4 regulates T-cell division, survival, and cytokine release (51). Overall, C6 carried genes with a predominantly inflammatory function.

#### Functional characteristics of microbial antigen responses

The C2 cluster was implicated in two of the dominant responses to microbial antigen-derived peptides, with the C2^hi^C4^hi^ and C2^hi^ types representing 33% and 14% of the responses, respectively. C2 consisted of 24 probes and was significantly associated with the GO term “immune response” with enrichment for this term driven by the immune-related genes CCL8, CXCL9, CXCL10, FCGR1A, FCGR1B, GBP1, GBP5, IFI44L, IFITM3, IFNG, and RSAD2. Many C2 genes carried interferon gamma (IFNG)-activated sequence at their promoters, suggesting C2 was linked to IFNG response. These genes included p65 guanosine 5’ triphosphatases GBP1 and GBP5, TF STAT1, cytokines CXCL9, CXCL10 as well as IFNG itself (52). GBP1 has antiviral effects including against influenza virus (53). STAT1 together with IFNG induce Th1 response by upregulation of TBX21 (54). IFNG is known to induce CXCR3 on lymphocytes, and CXCL9 and CXCL10 chemotaxis activated T-cells expressing CXCR3 to sites of infection or inflammation. Once recruited these cells stimulate local cells with IFNG to release more chemokines to further amplify inflammation (55). Expression of CCL8, RSAD2, and WARS, also in C2, can be induced by IFNG (56-58). To our knowledge, regulations of ANKRD22 and IFI44L genes by IFNG have not been investigated, but co-expression of these genes with other IFNG inducible genes indicates the possibility of induction by IFNG. C4, which together with C2 was high in SS3 was again significantly associated with GO term “immune system response.” This cluster consisted of 116 probes, and many of the genes in C4 enriched for this term were involved in antigen processing, which were also identified by biosets-analysis. These genes included CD74 which is involved in assembly and trafficking of class II molecules, subunits for class II molecules HLA-DMA,-DMB,-DPA1,-DQA1,-DQB1,-DRA,-DRB1,-DRB3, and-DRB4, immunoproteasome subunits PSMB8, PSMB9, and PSME1, and peptide transporter subunits TAP1 and TAP2. Genes in both C4 and the GO BP term also included IFN inducible genes and anti-viral genes such as IFI35, IFI6, IFIH1, IRF1, IRF8, OAS1, OAS2, and OAS3. Target genes of IRF1 include GBP1 for antiviral response, IL12 for Th1 response, and CASP1 for apoptosis (59). The OAS proteins produce 2’,5’-oligomers that activate RNaseL for RNA degradation, and degraded RNA can be a ligand for MDA5 and RIGI that induce IFN response during antiviral defense (60). Of note, other genes in C4 included IRF7, IRF9, and GTPases MX1 and MX2. IRF7 participates in a positive feedback mechanism during viral infection that results in enhanced expression of both IFNA and IFNB (59). MX1 and MX2 have been shown to have antiviral activities (61, 62). Overall C4 genes may promote antigen presentation and were involved in anti-viral responses. The other dominant responses to microbial antigens involved the pro-inflammatory program C3 described above and included the C0^lo^C3^hi^C6^hi^ type also encountered in T1D peptide responses (14% of responses to microbe-derived peptides) but also the C3^hi^ and C0^hi^C3^lo^ types. C0, which is described above, consists of an anti-inflammatory program; with C0^hi^C3^lo^ thus corresponding tentatively to an immunopressive response type that is the converse of the C0^lo^C3^hi^ response encountered in response to both T1D and microbial peptides.

Altogether, the unbiased definition of T-cell responses obtained employing a systems approach yielded a surprising number of types of responses given the relatively restricted range of antigen types used in this initial study. We found it possible to further refine response types with the definition of subtler sub-clusters as illustrated in **Figure6**. The heatmap focuses only on samples from SS2 with response type C0^lo^C3^hi^C6^hi^ and underscores the fact that more granular patterns of expression can be found within a given response type.

**Figure 6.**
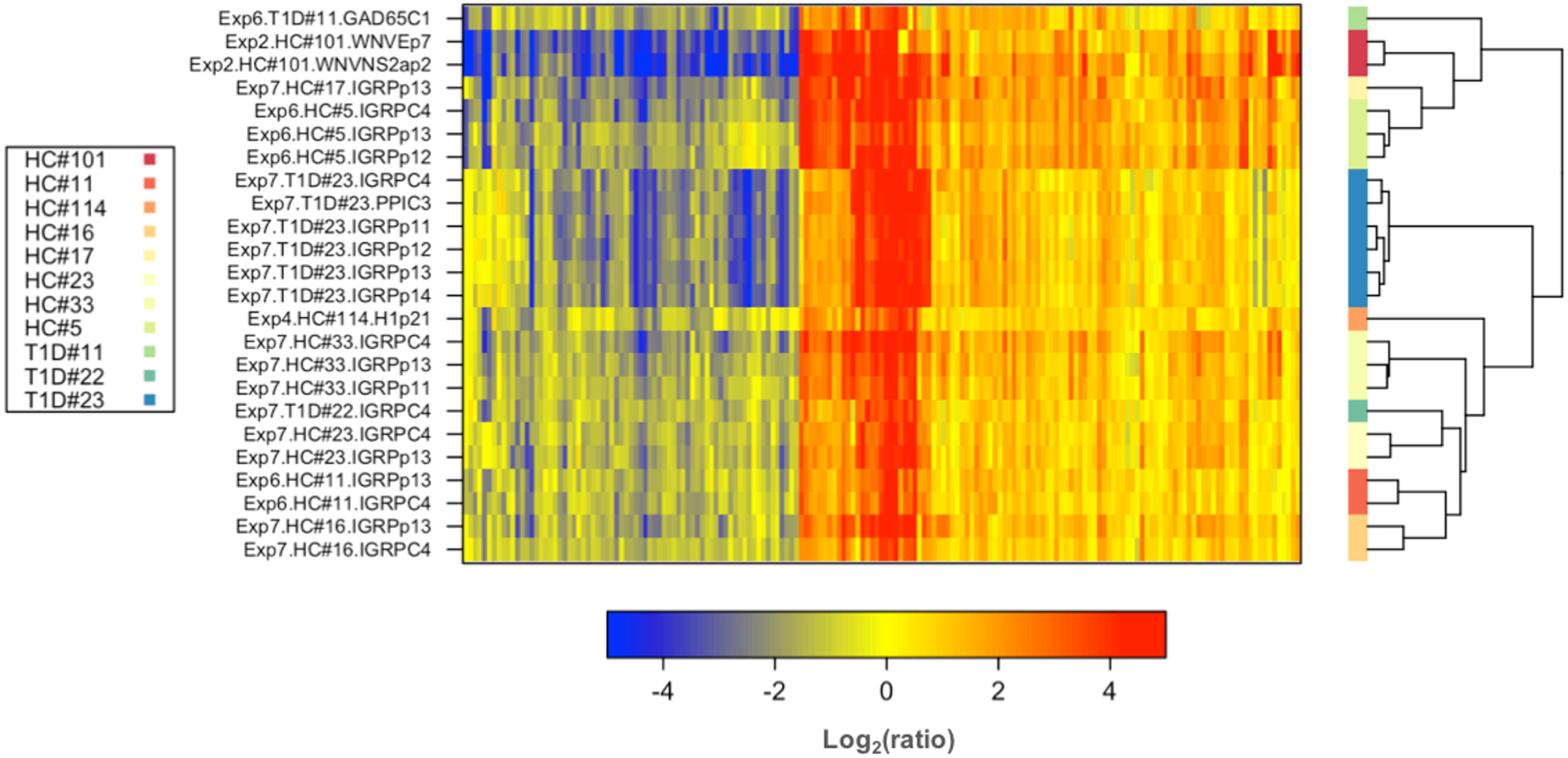
Sub-clusters can be found for a given type of response. This heatmap shows that distinct expression patterns can be found among samples constituting SS2 (rows) across genes C0, C3, and C6 gene clusters using (1-correlation) for the distance.

### Discussion

A compelling case can be made for the development of unbiased approaches for typing and characterizing antigenic T cell responses: Antigen-specific immunity plays a critical role in both health maintenance and pathogenesis. The successful introduction of therapies targeting checkpoints of T-cell immunity for treatment of cancer patients is only one of the most recent examples (63). Expanding the range of assays available for monitoring antigen-specific T cell responses is therefore likely to have a positive impact on development of new therapeutics and on clinical-decision making. Routine monitoring of antigen-specific immunity consists in the measurement of panels of cellular markers and cytokines released following peptide stimulation. The availability of so called “systems approaches” offers an opportunity to employ instead assays that simultaneously measure all the constitutive elements of a system (e.g. the PBMC transcriptome). This provides investigators with a global perspective on the molecular events occurring during antigenic responses. In addition, the approach has the distinct advantage of being unbiased, since it does not rely on *a priori* knowledge for panel selection. It simply measures everything that can be measured for that system.

The primary aim of the analysis presented here was to explore the potential utility of a systems-scale profiling readout for unbiased characterization and typing of antigen-specific T-cell responses. The method described in our paper suggests that this is indeed a viable approach, however it cannot at this stage be proposed as a new classification scheme. Indeed, the extensive collection of transcriptome profiles obtained following antigenic stimulations lacks diversity, being largely biased toward T1D peptides and included only a small number of non-T1D peptide-stimulated samples. Second, since the studies that have been assembled were carried out independently from one another, with different aims and sample sources, available “classical” immune phenotyping data are neither sufficiently uniform nor detailed to compare the classification scheme that we have obtained using a systems approach with the “gold standard” classification that uses cell surface markers and cytokine profiles accepted by the immunology research community. Nevertheless, what we have learned while performing these analyses should be useful to inform the design of future studies:

The whole genome transcriptional responses measured in over 300 samples stimulated with peptide *in vitro* were used for the stratification of the types of T-cell responses. On the basis that co-expressed genes likely share common regulatory mechanism and function for the same aim, we used cluster analysis to identify 9 co-expressed gene clusters and 11 sample sets. Interestingly, signatures combined in different ways to give rise to distinct types of responses. Given the the restricted antigenic repertoire used, the range of response types is likely to expand once responses to an extended range of microbes or allergens are measured.

To grasp the underlying biological functions of gene clusters, we examined them in detail using biosets-analysis and GO annotations (**Figure 3** and **Table 2**). C0 genes were enriched for a lipid metabolic process that possesses the ability to suppress T-cell responses. C2 genes were enriched for immune response and were IFNG inducible. C3 genes were enriched for pro-inflammatory genes. C4 genes were enriched for antigen presentation and anti-viral response. C5, C6, and C7 genes were enriched for defense response, inflammatory response, and immune response, respectively, that were both pro- and anti-inflammatory.

We also examined whether co-expressed gene clusters presented meaningful functional associations. SS3 was predicted to exhibit anti-viral response because it was enriched with Flu-MP stimulated samples (at a significance level of 0.05, data not shown). Statistical test revealed that C2 and C4, which carry IFNG inducible genes and antigen presentation and anti-viral response genes, were upregulated in SS3 (**Table 4**). Notably, several genes from these gene clusters, including IFNG, IRFs, MX1, OAS, and STAT1, are critical for controlling influenza virus infection (60, 64). Furthermore, Strutt *et al*. have shown that memory CD4^+^ T-cells promote anti-viral responses through production of IFNG, CXCL9, and CXCL10, and induce expression of MHC class II and CD40 molecules on CD11c+ cells (65). These genes also belong to C2 and C4, which were induced in SS3 in our study.

Sensitivity of the PBMC stimulation assay and readout was also evaluated. Organ-specific autoimmune diseases such as T1D are difficult to study because the target tissues are inaccessible. Easily available blood, carrying circulating immune cells, provides a great opportunity to examine immune status of an individual. However, the numbers of antigen-specific T-cells can be exceptionally small, hindering experiments that require large number of cells (66, 67). Using microarray, we found peptide-induced response from a million PBMCs that carried only 6 of flu peptide-specific CD4^+^ T-cells could be detected (data not shown). Because characterizing T-cells and monitoring their activities for progressive diseases are critical for understanding disease evolution, our method can be used to provide comprehensive qualitative immune responses. It should be noted that viably frozen PBMCs may not offer optimal condition for the identification of differential patterns of transcriptional responses. This was the case for instance, in experiments 6 and 7, where the number of peptide-specific CD4+ T-cells was expected to be low and responses of patients with T1D and healthy controls could indeed not be distinguished in our transcriptional assay. However, in a limited set of experiments, we have observed enhanced differential responses when comparing fresh vs viably frozen samples (data not shown). These preliminary data suggest that use of fresh cells would be indicated for the purpose of establishing a new T-cell classification scheme. Once a robust reference framework has been obtained it could be applied using a more sensitive targeted assay in clinical or research setting that necessitates the use of frozen samples.

Translation into a more practical and cost-effective assay can be achieved by reducing signatures to sets of representative genes. These genes in turn could act as surrogates for the entire set. The abundance of these transcriptional markers could be measured using conventional PCR or meso-scale profiling platforms such as high throughput qPCR or Nanostring (68, 69). It should be noted that the selection of a panel of analytes would be informed by a data-driven rather than a knowledge-driven approach. Such a strategy has already been implemented in the development of “transcriptome fingerprinting assays” that measure changes in blood transcript abundance *in vivo* (70).

Profiling the literature for transcriptional factors found among gene clusters identified in this study served to illustrate how systems-scale profiling can also contribute to identify potential knowledge gaps and to grow our knowledge of T-cell immunobiology (**Figure 4**). TFs and cytokines are drivers of immune responses. Some of these factors are known to play essential roles in T-cell development and functions. However, many others have never been examined.

Taken together, our findings demonstrate that transcriptome profiling can be used as a readout for measuring antigen-specific responses following peptide stimulation of PBMCs *in vitro*. Furthermore, identification of co-expressed genes allowed the development of a novel unbiased framework for the definition of types of antigen-specific CD4^+^ T cell responses. Thus, we believe the “systems approach” described herein can serve as a basis for further characterization of antigen-specific responses to expand current knowledge and to establish a foundation for a new generation of immunomonitoring assays.

## Acknowledgments

We thank all patients and volunteers who participated in this study, members of the Systems Immunology Division at BRI for their helpful feedbacks and uploading data to NCBI’s GEO. We wish to acknowledge investigators and staff of the BRI Translational Research Program and Clinical Core for subject recruitment, and for sample processing and handling. This work was supported by grants from the ITN (ITN N01 AI15416_)_, JDRF, and Baylor Health Care System and NIH contract HHSN272200900043C. The microarray data used in this study are available at NCBI GEO (series accession number: GSE72605).

## Supporting information captions

**S1 Table: Peptide sequence information.**

**S2 Table: Biosets gene list.**

**S3 Table: Gene cluster information.**

**S4 Table: Sample set information.**

